# A light-off response characterised by body contraction and ciliary arrest in *Acropora* coral larvae

**DOI:** 10.1101/2025.02.06.636794

**Authors:** Emelie A. Brodrick, Kei Jokura, Jamie Craggs, Rebecca Poon, Hannah Laeverenz-Schlogelhofer, Kirsty Y. Wan, Gáspár Jékely

## Abstract

Planulae of the reef-building coral *Acropora millepora* exhibit whole-body contraction and ciliary arrest after exposure to sudden light dimming. Quantitative behavioural assays confirmed reductions in swimming speed and alterations in vertical and horizontal movement under alternating light and dark conditions. High-speed microscopy revealed that ciliary activity ceases upon dimming, while the typically elongated body contracts into a rounded form, altering swimming behaviour. These larval responses suggest an adaptive mechanism to regulate positioning within the highly structured and complex light environment of reef habitats, critical for coral recruitment and survival.

## Introduction

The planula larva plays a critical role in the life cycle of a reef-building coral by dispersing from the parent colony. Within days of fertilisation and embryonic development, the larva must locate a suitable habitat to attach to, where it will settle and metamorphose into the primary polyp of a new coral colony. The animal will spend the rest of its life at that location, filter-feeding with tentacles (primarily at night) and utilising organic products of algal endosymbiont photosynthesis accrued during the daytime (Sorokin, 1973). Coral colonies can subsequently increase in size through asexual budding, producing clonal polyps.

Corals (Anthozoa, Hexacorallia, Scleractinia) are cnidarians, the sister group to all bilaterians (Genikhovich and Technau, 2017; Hejnol et al., 2009). Their nervous systems, in both larvae and adults, are not centralized into a brain-like structure; rather they consist of nerve nets distributed across radially symmetric bodies (Satterlie, 2019). Planula larvae have a nerve plexus at the anterior aboral pole, which usually underlies an apical organ of sensory cells. Despite the relative simplicity and diffuse, non-centralised organisation of their nervous systems,coral larvae can make decisions and are known to actively explore the substrate, ndefinedchoosy in where they settle (Harrington et al., 2004; Maida et al., 1994; Ricardo et al., 2017). They have been shown to use a multitude of sensory cues to inform their decision for successful settlement (reviewed in (Pysanczyn et al., 2023)).

Light availability in clear, relatively shallow water is an essential requirement for the successful recruitment of corals hosting photosynthetic endosymbionts, and tropical coral reef communities often display disctinct depth-related zonation(Mundy and Babcock, 1998; Roberts et al., 2019). The staghorn coral *Acropora millepora* produces relatively large (500–900 µm), lipid-rich larvae that swim slowly and do not acquire endosymbionts until after metamorphosis into a polyp (Quigley et al., 2018). Anthozoan cnidarians lack visual eyes or recognisable ocular structures at any life stage. Nevertheless, light intensity and spectral composition are well-established cues influencing larval settlement behaviour (Foster and Gilmour, 2016; Gleason et al., 2006; Mason et al., 2011; Mundy and Babcock, 1998; Ricardo et al., 2021; Strader et al., 2015). Consequently, larval light sensing is crucial for the survival and ecological success of many reef-building coral species.

Adult *A. millepora* corals sense light using non-visual cryptochromes (Gorbunov and Falkowski, 2002; Levy et al., 2007) and opsins (Gornik et al., 2021; Vize, 2009) to monitor solar and lunar cycles, enabling the timing of nocturnal feeding and the synchronisation of annual spawning events (Boch et al., 2011). The motile planula larvae of many scleractinian corals are also responsive to light. At least six genes encoding opsin (Acropsin) apoproteins have been identified in *A. millepora* larvae, which can form light-sensitive photopigments when coupled to a retinal chromophore (Mason et al., 2012; Mason et al., 2023). However, these opsins are not localised to specialised photoreceptor cells with distinctive ocular morphology, such as extensive membrane elaborations or shading pigments. Instead, opsin-expressing cells are scattered across the larval ectoderm and closely resemble neighbouring epithelial cells (Mason et al., 2012), making them difficult to identify using morphological approaches such as electron microscopy.

In accordance with the expression of several opsins (Mason et al., 2023), coral larvae exhibit light-regulated behavioural responses including photomovement (Mulla et al., 2021) or putative phototaxis in some species (Kawaguti, 1941; Lewis, 1974), and spectral preferences during settlement (Foster and Gilmour, 2016; Mason et al., 2011; Mundy and Babcock, 1998). Notably, Sakai and colleagues (Sakai et al., 2020) demonstrated reductions in average swimming speed within batches of *A. tenuis* larvae that were roughly proportional to the white light intensity-decrease that they were exposed to. They further tested the response with different wavelengths of light, showing that swimming speed is most affected by changes in blue light intensity, while not responsive to changes in red light. Here, we examine responses of *A. millepora* larvae to light dimming in further detail. We confirmed the presence of this light-off response in this species and also found that the coral larvae undergo whole-body muscular contraction in addition to ciliary arrest after a sudden dimming in light intensity. We propose a simple model for position maintenance in the water column and on the reef, in the absence of spatial vision.

## Materials and Methods

Colonies of *ex situ A. millepora* synchronously spawned (1 Dec 2021 & 14 Nov 2022) at the Horniman Museum and Gardens Aquarium facility, London, UK. The gametes from different colonies were cross-fertilised to produce embryos, which were transported to University of Exeter, UK when they were 3 days post fertilisation (dpf) and our experiments were carried out between 5 and 7 dpf. Larvae were stored in 500 ml covered glass containers in seawater at 27 °C inside an incubator, with aquarium lighting (Marine 3.0 Nano, Fluval, Quebec, Canada) providing a 12/12 hr light/dark cycle.

Unless specified, all experiments were carried out in a 27 °C temperature-controlled room to prevent temperature fluctuations and all experiments were conducted in a dark room with no external sources of visible light. Before experiments, larvae were acclimatized to the light-on conditions for at least 5 minutes. This allowed the first light-on period in experiments to serve as a proxy for baseline behaviour in constant light conditions.

Light spectra were measured from each experimental apparatus (Fig. S1 A-D) from the position that larvae were held, in absolute irradiance using a calibrated spectrophotometer (QE65000, Ocean Optics, Orlando, USA) coupled to an optical fibre and cosine corrector. The total absolute irradiance from a light source was then attained by calculating the area under the spectrum curve.

### Free-swimming horizontal assays

Larvae (10-15 individuals, *n=*6 batches) were placed in a 5 mm shallow glass cuvette (20 x 9 mm) containing 900 µl seawater and allowed to swim freely (apparatus depicted in Fig. 1A). They were recorded using an infrared-sensitive USB camera (iDS UI-3360CP-M-GL Rev.2, Imaging Development Systems, Obersulm, Germany) mounted on a stereomicroscope (Leica M205, Wetzlar, Germany). An infrared ring light (LED-HP6095-3, LeeTun Optical Tools, China) with peak emission at 850 nm wavelength was positioned below to create dark-field illumination, with absolute irradiance of 2543.7 µW/cm^2^/nm. A 780 nm long-pass colour filter (FGL780S, Thorlabs, Newton, NJ, USA) was placed in front of the camera to stabilize the image during light stimulus changes.

**Fig. 1.**
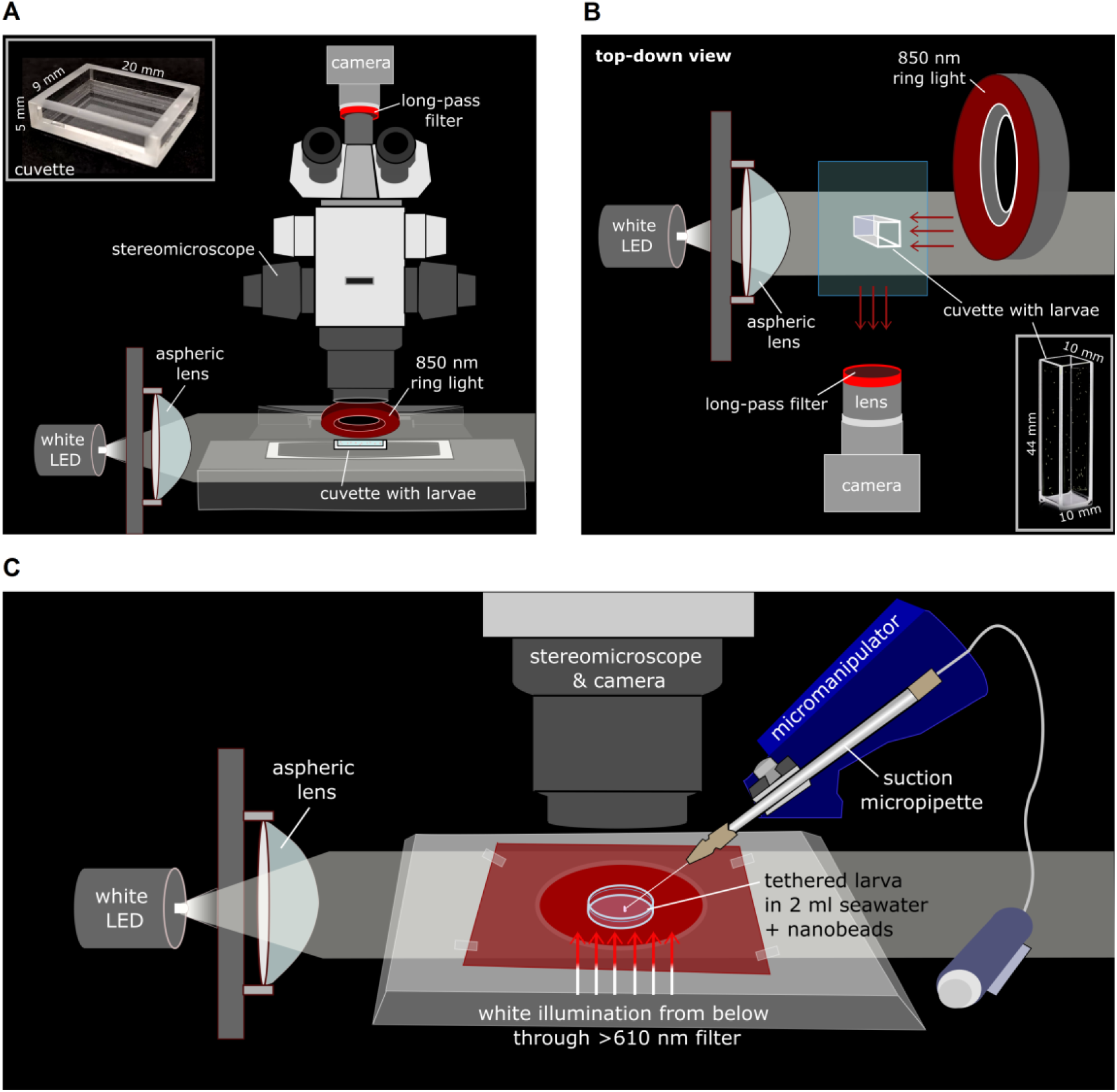
Schematics of the apparatus used to assess photobehaviour. **A)** Free-swimming larvae held in a shallow cuvette to assess horizontal movement in response to white light on/off stimulation. **B)** A single larva is tethered on a suction micropipette and held in a dish of seawater containing nano-particle beads that allow visualization of flow fields and measurement of bodily shape changes. **C)** Top-down view of apparatus used to assess vertical movements of free-swimming larvae inside a 44 mm high cuvette.

Light stimulation was provided from one side (which alternated) during the first 90 s using a white LED (MCWHL7, Thorlabs, Newton, NJ, USA) controlled with an LED driver (LEDD1B, Thorlabs, Newton, NJ, USA). The beam was passed through a collimating lens (LAG-65.0–53.0 C with MgF2 Coating, CVI Melles Griot) to produce a bright (5 cm diameter) spot of light with uniform intensity to fill the whole area of the cuvette. Larvae were first acclimatized to the bright light for 5 minutes, with total absolute irradiance of 1421.6 µW/cm^2^/nm. Starting with the stimulation light on, 90-second alternating photoperiods (light on/off) were shown twice, making a total experiment time of 6 minutes. Experiments were carried out in an otherwise dark room so that there was no visible light available to the larvae during the light-off periods, resulting in measured absolute irradiance 0.0 µW/cm^2^/nm (spectral data in Supplementary Fig. S1A).

Video data from six replicate experiments were analysed with a custom Fiji macro, which used the Mtrack2 plugin (Stuurman, 2003, https://valelab4.ucsf.edu/~nstuurman/IJplugins/MTrack2.html). The video was sampled into 5 sec (100 frame) clips and from each clip, xy coordinates of thresholded larval profiles were extracted between video frames. Custom scripts in R (version 4.1.3 (R Core Team, 2021)) were used to analyse the data, then calculate and plot swimming speed, in addition to swimming trajectories with respect to the direction of the light source. To allow for adjustments in behaviour to ensue, speed and horizontal displacement data (distance travelled in x per time point) from the second 45 s half of each time period were compared statistically with Mann-Whitney Wilcoxon rank sum tests, due to non-normality within the datasets (Shapiro-Wilk tests, p<0.05). All analysis code, source data, and video data supporting this manuscript are available online at (Brodrick et al., 2025).

### Body shape changes

Data were collected over two years, 2021 (6 dpf, n = 6) and 2022 (6 dpf, n = 6). One at a time, larvae were held in a 30 mm diameter Petri dish of filtered seawater under a Leica M205 stereomicroscope. A custom-pulled suction micropipette (TW-1200 borosilicate glass) attached to a micromanipulator was used to tether larvae from the lateral body wall or near the oral pole. White light from below the dish was passed through a red long-pass filter (>610 nm), providing constant background illumination for imaging. Red light was used as the “dark” phase rather than near-infrared so that nanobeads could be visualized effectively for assessment of ciliary flow fields (see next section). Video data were collected through a 620 nm long-pass colour filter using a USB camera (iDS UI-3360CP-M-GL Rev.2) with near-infrared sensitivity (apparatus shown in Fig. 1B).

A cool-white mounted LED provided unilateral illumination. The beam was passed through a collimating lens (as above) to ensure directional stimulation from one side. Following tethering, each larva was allowed to acclimatize for 5 min under LED illumination. The recorded trial began after this acclimatization period. The experiment consisted of repeated light-on/light-off transitions. The LED-on periods served as the adapted baseline state, whereas the transition to LED-off constituted the experimental stimulus. In Year 1 (2021), larvae experienced three alternating light-on/light-off cycles. Each cycle lasted 90 s, except for the first light-on period, of which only 30 s was recorded. In Year 2 (2022), larvae experienced two full 90 s light-on/light-off cycles. Light intensity and spectral composition were kept constant between years (Supplementary Fig. S1B).

Larval profiles were measured from thresholded images with standard built-in ImageJ functions and plugins. Body shape metrics were quantified at 10 s intervals throughout each recording. However, to avoid transient effects associated with illumination transitions, only the second half (final 45 s) of each 90 s light condition period was included in statistical comparisons. As each larva experienced two light on–off cycles, values from the analysed time windows were averaged within each illumination condition for each individual prior to statistical testing, such that the larva constituted the unit of replication. Larvae that were weakly tethered and rotated or tilted substantially during recording were excluded from analysis.

### Ciliary flowfields

The 6-minute videos from the 2022 dataset (5 dpf, *n*=6) used to measure body shape changes were additionally analysed to assess flow fields created in the surrounding water by the larva’s beating cilia (Fig. 1B). The seawater contained 1 µm Fluoresbrite® multifluorescent microspheres (volume fraction 10−5 v/v) to allow measurement of the flow fields and the video was captured at 90 fps. The videos were analysed with the PIVlab tooblox (v2.63) in MATLAB (2023b, MathWorks, Natick, MA, USA). After generating flow vectors, the mean velocity magnitude was measured from regions of fluid flanking the side of each larva between frames of the videos. The location of these areas were adjusted at times during the video to allow for larval shape changes and movement of the larva’s position on the micropipette. The raw data were smoothed by averaging across each second (90 frames). Larvae that were held weakly and moved considerably on the pipette (rotated or tilted) were excluded from analysis.

### Ciliary beat frequency

Larvae (5 dpf, *n=*2) were tethered with a custom-pulled TW-1200 borosilicate glass suction micropipette (90/30 µm OD) attached to a micromanipulator and held in a dish of seawater on a Leica DMi8 inverted microscope stage. Imaging was conducted at 22 °C room temperature (because the microscope could not be moved to the 27 °C room). Video data were collected from the microscope in Differential Interference Contrast (DIC) with 40x objective (Leica HC PL FLUOTAR L 40x/0,60 CORR PH2) and a Phantom V1212 camera. Regions of ectodermal locomotor cilia were recorded at 500 fps from previously light-adapted larvae. The larvae were imaged under bright white light for about 30 s, then the light was switched to dim red-filtered (>600 nm) light for the remainder of the recording (spectral data in Supplementary Fig. S1C). The camera gain was increased by 4.5x to adjust for the lowered light intensity.

The ciliary zone within each frame was located by thresholding the image to find the boundary of the (opaque) larval body. The cilia project approximately 20 µm outwards from the body surface. For each frame, multiple small boxes were drawn within the ciliary band, parameterised by the arc length, *d.* The boxes extended 40 pixels (14.4 μm) in the direction normal to the larval body surface, and 13 pixels (1.8 μm) in the *d* direction. Within each box, the standard deviation of the image intensity was calculated for each video frame, giving a time series at each location *d*. A wavelet transform was then calculated from the time series at each value of *d* to give the time-resolved frequency spectrum of the signal at that location. The magnitude of the wavelet transform was averaged over the 6-12 Hz frequency band, since the periodic beating occurs at a frequency of approximately 8-10 Hz. This value indicates how periodic the signal is as a function of time, thus giving a high value when the cilia are beating, and a low one when they are stationary. These averaged wavelet transforms of all of the locations are stacked to produce kymographs.

### Vertical positioning

Free-swimming larvae (5 dpf, *∼*60 individuals, *n=*6 batches) were placed into a glass cuvette of seawater measuring 44 mm height (10 x 10 mm width and length) and illuminated from one side with the near-IR ring light (apparatus depicted in Fig. 1C). The same infrared-sensitive camera with lens and long-pass (780 nm) filter used in previous experiments captured video (20 fps) from directly in front of the cuvette. On the opposite side of the IR ring light, a cool-white LED passed through a collimating lens provided an even beam of light stimulation in 90-second on/off photoperiods that repeated twice (6 minutes total). Light intensity values were kept the same as in the horizontal swimming experiment (spectral data in Supplementary Fig. S1D).

Video data were analysed to track larval positions using a similar approach to that described for the horizontal free-swimming assay in shallow cuvettes. Shorter tracking intervals of 1 s (20 frames at 20 fps) were used to accommodate the higher density of larvae and the increased frequency of track overlaps in the vertical cuvette, thereby reducing loss of positional information during tracking.

To assess the natural buoyancy of the larvae, the cilia were removed temporarily by exposing larvae to hyperosmotic seawater (3% NaCl in seawater) for around 30 s until swimming movements ceased. Larvae were washed in clean seawater immediately after. This deciliation method was reproduced from Takeda-Sakazume and colleagues (Takeda-Sakazume et al., 2022). The deciliated larvae were placed inside the vertical glass cuvette and recorded (with light stimulus on) on the camera (20 fps). The water was mixed by pipetting water up and down twice to redistribute the larvae in the water column. Then, trajectories of the larvae were recorded for 50 s as they moved inside the cuvette according to their buoyancy. The deciliated larvae were kept overnight to confirm that they were alive and healthy during the experiment. Observations the next morning confirmed not only survival, but that the larvae had regrown ectodermal cilia and were once more swimming as usual.

Statistical analyses were performed by comparing speed and vertical displacement data (distance travelled in y per time point) from the middle 30 s of each dark period with the middle 30 s of the second light period. Mann-Whitney Wilcoxon rank sum tests were performed due to non-normality of the data sets (Anderson-Darling tests, p<0.05). All experiments had a trial-based design based on the signal detection theory (Green and Swets, 1966). We measured response to repeating events of stimulation (light off) separated by baseline light on periods. This trial-based approach allows us to distinguish the response of the larvae from the noise.

## Results

### Horizontal swimming speed is reduced during dark periods

To assess whether *A. millepora* larvae display a similar light-off swimming speed response to that of *A. tenuis,* (reported by (Sakai et al., 2020)), free-swimming larvae (6 batches of 10-15 individuals, *n*=109) were tracked inside a shallow cuvette. Horizontal movement speed was assessed while the light stimulation conditions were alternated between 90 s white light (absolute irradiance 121.6 µW/cm^2^/nm) and 90 s darkness (0.0 µW/cm^2^/nm).

Under the initial phase of bright white light, the pre-light-adapted larvae swam at a mean speed ± its standard deviation of 0.41 ±0.43 mm s⁻¹, with maximum speed of 2.0 mm s⁻¹. Mean swimming speed was altered by the light changes (Fig. 2A, see Video S1 for example) and comparing the final 45 s (second half) of bright time periods with dark periods revealed a significant reduction in swimming speed in the dark (Wilcoxon rank sum test *W*=183558, p<0.001, Table S1 for full statistical test results and Fig. S2 for individual batch figures). Larval swimming speed reduced around 10 s after the onset of darkness, and during the final 60 s of the dark phase, it averaged 0.10 ±0.16 mm s⁻¹. In the following light phase, swimming recovered gradually until it reached a speed of 0.51 ±0.45 mm s⁻¹ in the final 30 s of this light phase. Finally, larvae in the second dark phase showed a similar response to the first, with relatively rapid reduction of swimming speed (within 10-20 s), reaching a mean of 0.14 ±0.23 mm s⁻¹ in the final 30 s.

**Figure 2.**
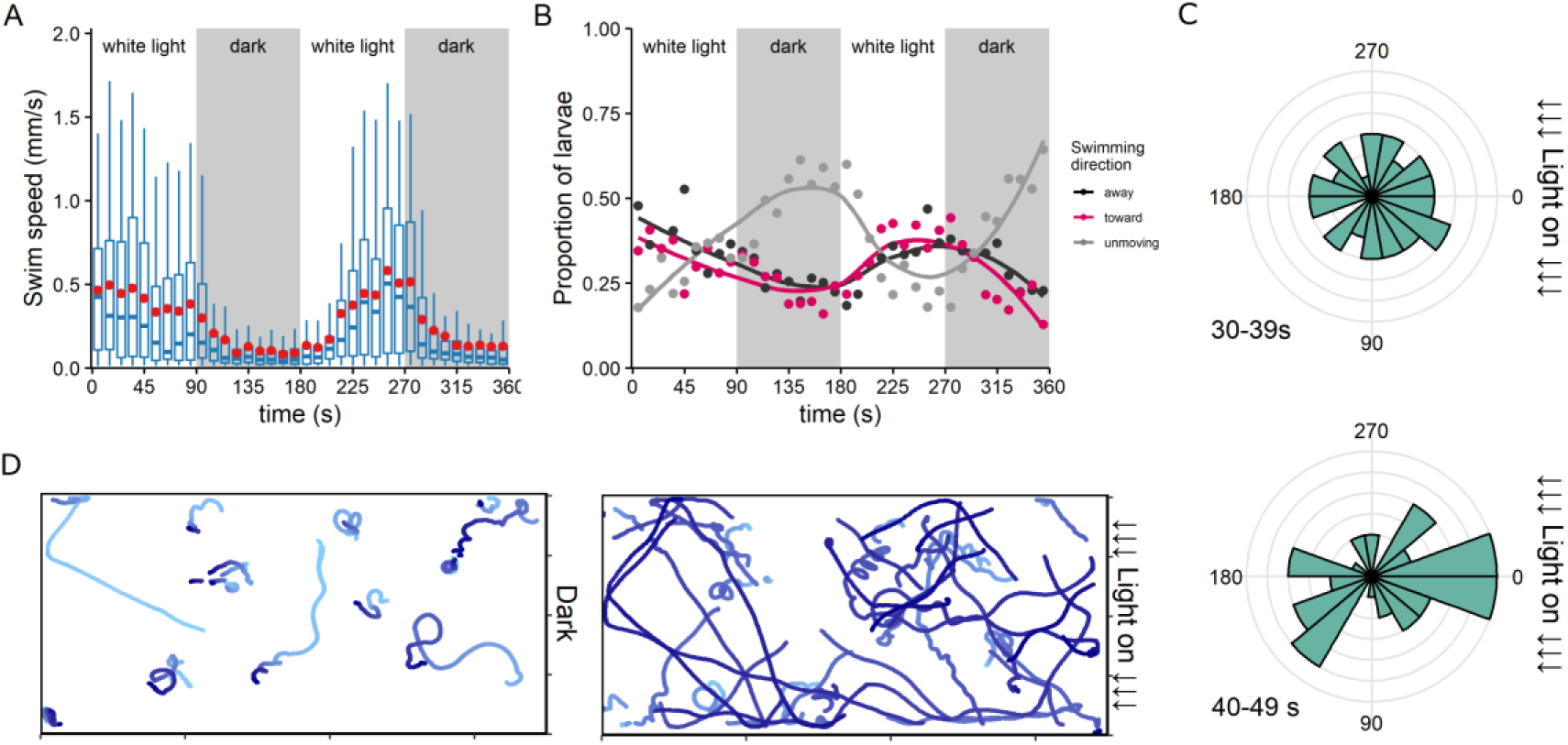
Free-swimming photobehaviour of A. millepora larvae in shallow cuvettes. Light stimulation was provided in alternating 90 s intervals of collimated bright white light and darkness. Tracking of individual larval positions allowed measurement of (A) swimming speed represented as boxplots, with red points overlaid to show mean values. Boxes represent interquartile range with median centre line; whiskers extend to the most extreme values within 1.5 × the interquartile range. (B) Swimming direction as a proportion of the total number of larvae, with respect to position of the light source, averaged across six batches of 10-15 larvae. C) The angle of larval trajectories over 10 s for all six batches are shown as circular histograms of 20-degree bin width. The data come from the second light-on period, 30-39 s and 40-49 s after the stimulus light came on. Those travelling at zero degrees (right) are moving directly towards the light source. Grey lines represent larval counts of 1. D) Example batch showing trajectories (x,y) of larvae over time in the first dark period (91-180 s) and following light-on period (181-270 s). Increasing time is represented by increasing darkness of the blue shade of the points. Dimensions of cuvette area are 20×9 mm.

### Larvae showed no evidence of phototaxis

Swimming direction in a horizontal plane was not influenced by the position of the light source (Fig. 2B) and the proportion of larvae swimming towards or away from the light source remained equal to one another during the bright light periods. The proportion of larvae considered to be “unmoving” (travelling less than 0.1 mm s⁻¹ in any direction) increased (to >0.5 of the total) during the second half of the dark periods. Comparing horizontal displacement in the second half (last 45 s) of each light condition, we found no difference between bright light and dark (Wilcoxon rank sum test *W*=338476, p=0.273, Table S2 for full statistical test results and Fig. S3 for individual batch figures).

Figure 2C shows the angles of larval trajectory over 10 s after the light stimulus came on the second time. We show here 30-39 s and 40-49 s after stimulus change because that is when most larvae began to actively swim and travel again in the cuvette after spinning on the spot during the dark phase. In both periods, larvae travelled in random directions with respect to the light source at 0°, especially between 30-39 s. It should be noted that due to the rectangular shape of the cuvette, it would be expected that there would be greater movement in the horizontal plane (0° and 180°), as is seen between 40-49 s. This shows that when light stimulation is presented after a dark period, larvae do not swim towards the light source. To visualise the difference in swimming behaviours, larval trajectories for one example batch (#2) are plotted for the entire first dark period and the following light period (Fig. 2D). Supplemental Video S1 shows the movement trajectories for another batch (#3). The larvae tended to spin on their own axis in the dark, as shown by the small loops. In the following light-on phase, greater distances are covered in random directions.

### Ciliary arrest and flowfields

Individual tethered larvae (*n*=6) were also examined more closely while ambient light conditions alternated between bright white light and dim red light. Using particle image velocimetry, we measured the flow of water movement from an area flanking the side of the larvae, created by the beating cilia of the larvae (Fig. 3, see video S2 for example of raw data). This allowed an indirect assessment of ciliary swimming activity.

**Figure 3.**
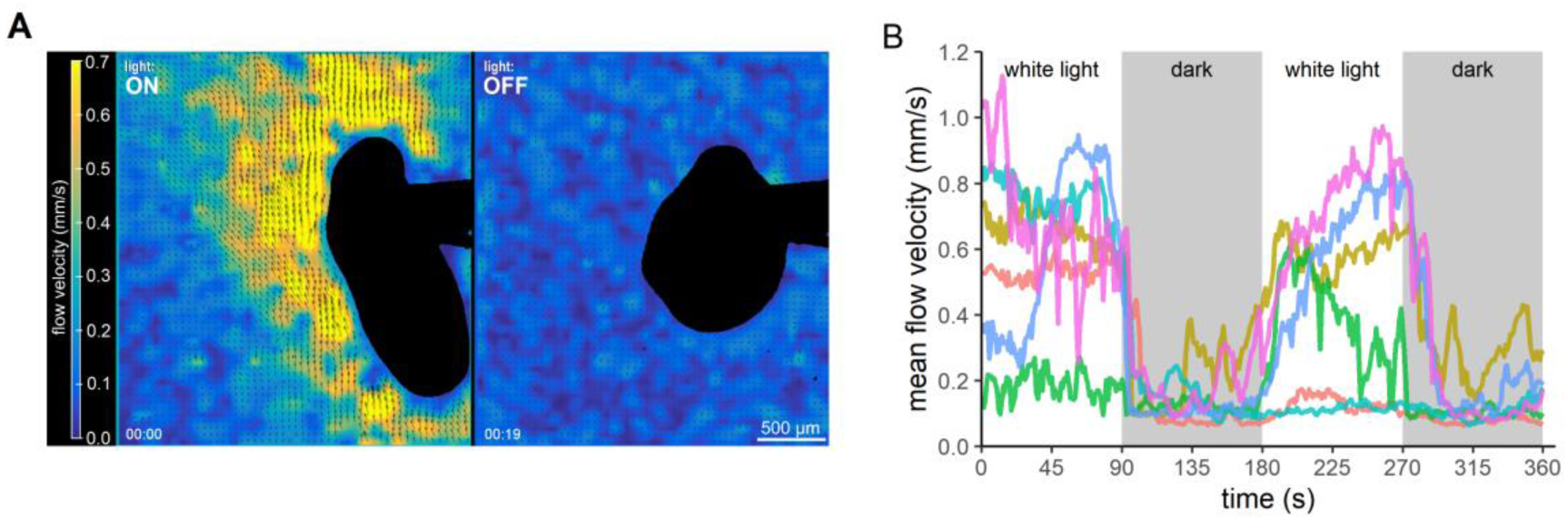
Flow velocity (mm s⁻¹) of water flanking tethered larvae. **A)** Examples of PIV analysis showing active flow with white stimulus light on (left) and then 20 s later (right) when the stimulus light had been off for 15 s. **B)** Flow velocities for six individual larvae are represented in different colours over time as light intensity was alternated between bright white light and dim red light. The larvae create these flows with their swimming cilia and they are measured by particle image velocimetry using nanobeads in the water.

Overall, fast active flow was recorded around larvae during light-on periods, then flow reduced or stopped completely during the dark phases (Fig. 3A & B, examples in Videos S2 & S3). There was a considerable range in adjacent water flow-speed between individuals under bright light conditions (0.5 - 1.0 mm s⁻¹) and flow speed fluctuated considerably over short timescale during light periods. One larva was not swimming actively during the first light phase, and two other larvae did not resume fast swimming in the second light-on phase. However, the flow reduction response to the decrease in white light intensity was consistent. The flow reduced to its minimum within 15 s after white light dimming (e.g. Fig. 3A) and in most cases sustained a very low speed until the following light phase. Similar to the trend in swimming speed, the recovery of fast flow in the second light phase was gradual compared to the light-off response (Fig. 3B).

To directly evidence ciliary arrest in response to change from bright white light to dim red light, we performed high-speed imaging at 500 fps of ectodermal swimming cilia from several larvae. Here, we illustrate two examples (Fig. 4), which show that although a ciliary arrest happens for both, the time between the change to dim red light and the stop in ciliary beating varies quite considerably, from around 4-43 s. Our first example (Larva #1), where an aboral pole region was imaged (Fig. 4Ai), the cilia show (slightly patchy) beating during the initial 30 s in bright light (Fig. 4Aii). Just 4 s after the switch to dim red light, all cilia stop their beating simultaneously, demonstrating a rapid light-off response (Fig. 4Ci, Video S4). After around 20 s of the red light condition, the cilia from *d*=200-370 µm recovered and displayed intermittent beating, whilst the rest of the cilia remained stationary for the rest of the imaging period (Fig. 4Aii).

**Figure 4.**
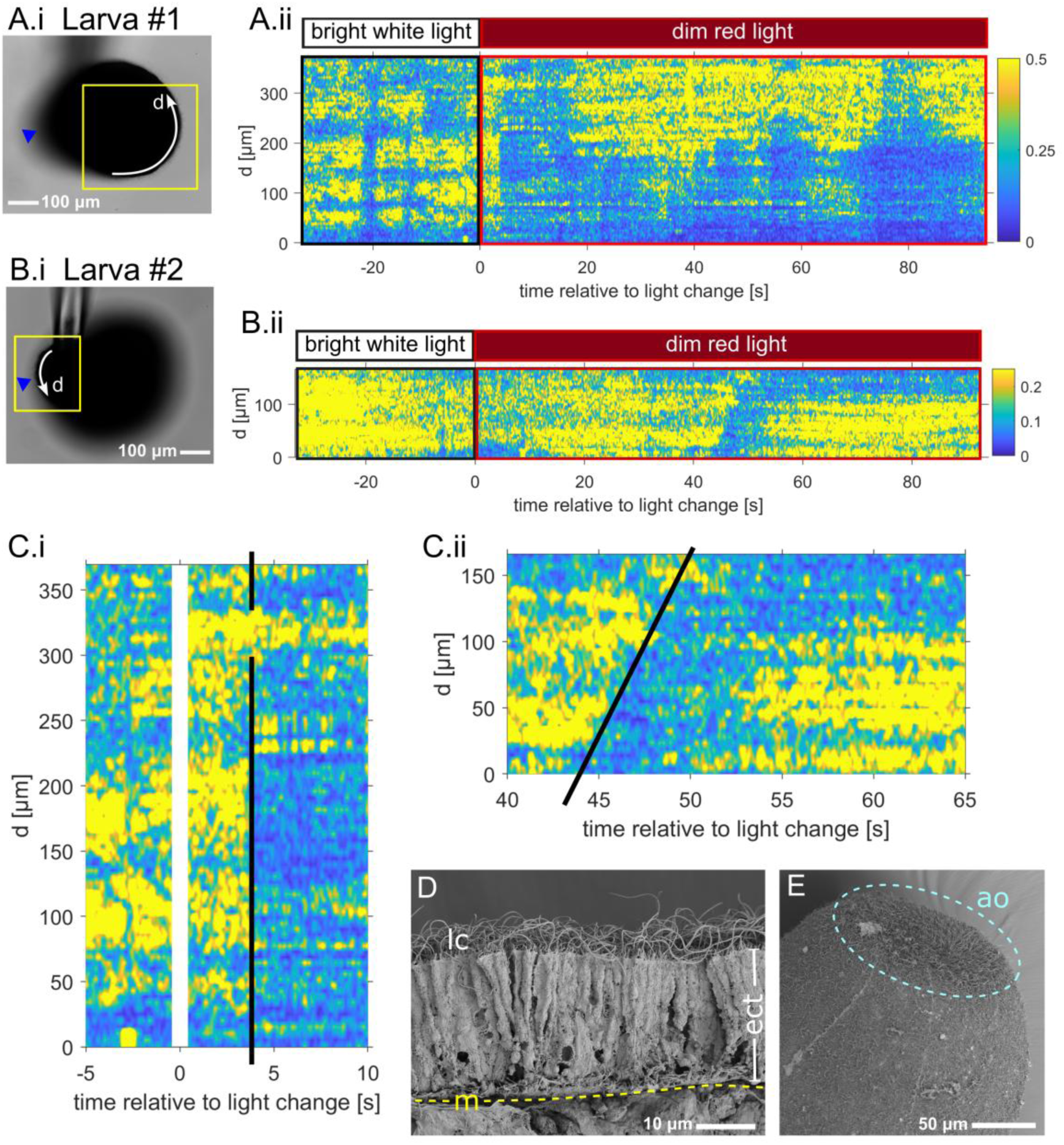
Ciliary beating responses of two individual larvae to light dimming revealed by high speed imaging. Larva #1 (**Ai**) and Larva #2 (**Bi**) viewed with low magnification, tethered on the micropipette. Yellow boxes indicate the region of interest from which high-speed videos were recorded. Blue arrows show the location of the oral pores, so that Larva #1 shows data taken at the aboral/anterior end of the body, and Larva #2 at the oral/posterior end. The data are spatially resolved so within each video, different locations of the cilia around the larval body are parameterised by the arc length d, indicated by the white curved arrows. This region was limited to the section of the ciliary surface that was in focus. **Aii & Bii)** Kymographs showing the magnitude of the periodic signal from the ciliary surface of Larva #1 (Aii) and Larva #2 (Bii) during a light-off response experiment. The colour indicates strength of the periodic signal in the video as a function of arc length (d) along the edge of the larva and time (s). A high value (yellow) means that the cilia are beating periodically, whilst a low value (blue) shows that the cilia are not beating. **C)** The ciliary arrest response is shown in more detail, shortly (4 s) after the light dimming for larva #1 (i) and more than 40 s after the dimming for Larva #2 (ii). A black line indicates the pattern of arrest in space, d, along the edge of the larva. **D)**SEM image of the fractured ectoderm (ect) of a fixed A. millepora larva showing the dense locomotor cilia (lc) protruding from each long, narrow ectodermal cell. Mesoglea (m, yellow dashed line) separating the two cell layers is shown for reference. **E)** SEM image of the aboral pole of a larva and its apical organ (ao, inside dashed ring). Note the different appearance of the ciliated ectodermal surface.

Larva #2 was imaged nearer the oral pole (Fig. 4Bi) and the strength of the ciliary beat frequency periodicity was weaker here (0.2, compared to 0.5 at the aboral region of larva #1). The cilia here were slower to respond after the change to dim red light (Fig. 4Bii, Video S5). The cilia at *d*=0 µm stopped beating first, 43 s after the stimulus light dimming. Then, the arrest spread progressively across the ectoderm at a rate of 23.3 μm s⁻¹. It reached *d*=166 µm around 7 s later, producing the sloped boundary seen in Fig. 4Cii. Soon after all the cilia had stopped, those within the *d*=0-100 μm region recovered their beating, whereas the other section of cilia (*d*=100-166 μm) remained motionless. These two examples demonstrate that there is high temporal variability and spatial patchiness within the ciliary response.

To assess whether the uneven ciliary arrest followed a consistent spatial pattern, such as initiation at the oral pole and propagation towards the aboral pole, we performed additional particle image velocimetry analysis using smaller spatial sampling regions. Five regions were defined along one side of the larva, spanning from the aboral (anterior) to oral (posterior) poles, and flow was analysed over the 50 s period immediately following light dimming (Fig. S4). Across individuals, flow velocity decreased rapidly after light-off in all measured regions, typically within the first 5–15 s, indicating a prompt reduction in effective ciliary propulsion. However, both the magnitude and timing of this reduction varied substantially between larvae and across body regions. Oral regions consistently exhibited lower flow velocities than aboral and lateral regions, both before and after light dimming. Rather than a uniform or simultaneous arrest, reductions in flow were spatially heterogeneous, and in several larvae, gentle flow persisted or re-emerged transiently following the initial decrease. Together, these observations indicate that ciliary arrest during the light-off response is partial and variable, rather than reflecting a tightly coordinated, all-or-none motor response.

### Body shape changes

Individual larvae of the same age, developmental stage and batch can vary considerably in overall size, shape and eccentricity. Generally by 5-6 dpf, the body has elongated to a bullet-shape, often with slightly wider profile at the aboral (anterior) pole.

All larvae (*n*=12) demonstrated dark-induced shape changes during the light-off experiment, by gradually contracting the longitudinal muscles and shortening their body length along the oral-aboral axis. This resulted in a reduction of length/width ratio. This shortening resulted in a rounder shape with some widening across the middle of the larvae with axis perpendicular to the oral-aboral axis. This significantly changed the eccentricity (width to length aspect ratio) of the larvae (Wilcoxon rank sum test, *W*=1059, p=0.001 for 2021, *W*=551, p<0.001 for 2022) (Fig. 5A) and also, to a lesser extent, the 2-dimensional area of the larval profiles (Wilcoxon rank sum test, *W*=1262, p=0.04 for 2021, *W*=716, p<0.001 for 2022) (Fig. 5B) (statistics in Table S3). Generally, the response showed good reproducibility and in most larvae, the change was striking (see example in Fig. 5C). However, one larva from 2021 spawning showed a very weak response to the first and third dark periods, suggesting that the response may not occur consistently. The two cohorts of larvae measured generally within the same size range, although larvae spawned in 2022 displayed greater changes than the previous year. The response can be watched in Video S2 and Video S6, where two larvae (one from each cohort) are exemplified.

**Figure 5.**
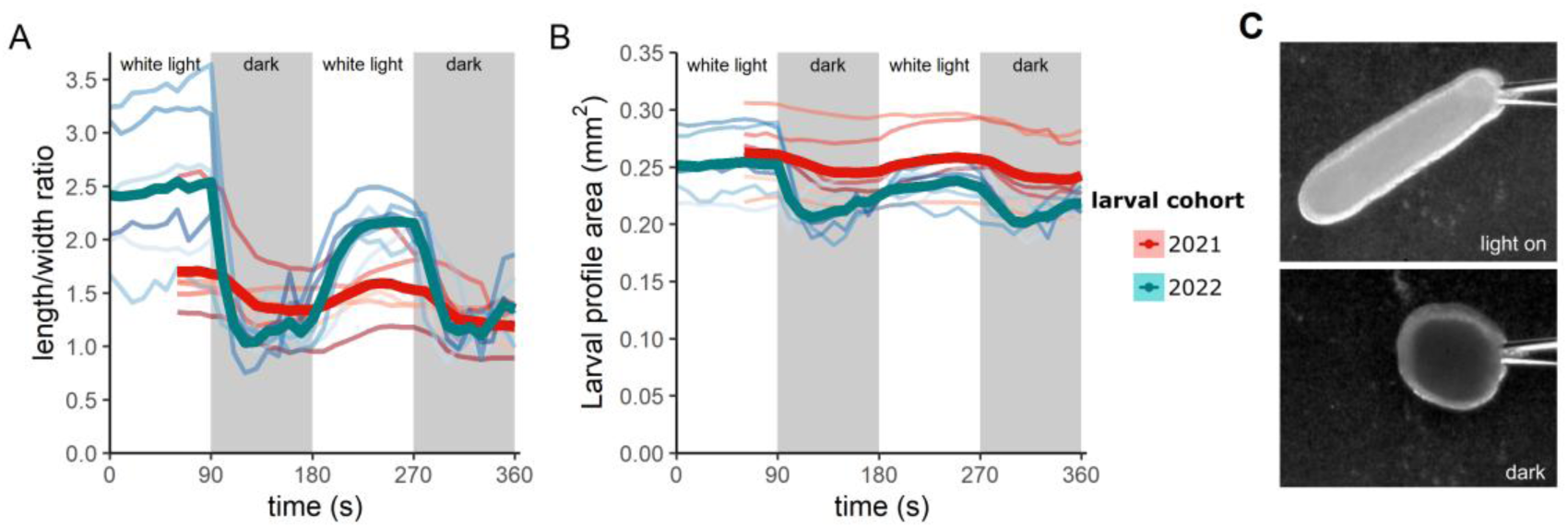
Larval body contractions and shape change in response to decrease in white light intensity. Data are shown for individuals measured in 2021 (red shades) and 2022 (blue shades). The means per cohort are represented in a stronger shade and thicker line. 90 sec bright white light periods were alternated with 90 sec dark periods, which are indicated by grey background shading. Note, individuals from 2021 were not recorded for the first 60 s of the first light period. **A)** Eccentricity changes as a ratio of length (along oral-aboral axis) divided by width (perpendicular to the oral-aboral axis) are represented over time. **B)** The projected area of the larval profiles as they appear in the video. **C)** An example of a larva held on a suction micropipette near its oral opening, elongated during exposure to bright white light at the beginning of the experiment (top). The same larva is shown again after 60 seconds into a dark phase (bottom) and its body is contracted in length to form a spherical shape.

### Vertical positioning

To assess the natural buoyancy of larvae, we removed the cilia using a brief exposure to hyperosmotic seawater. This made them unable to swim and actively modulate their own position in the water column. After mixing to distribute the larvae inside a tall (44 mm) and narrow (10 x 10 mm) glass cuvette filled with artificial seawater, almost all larvae floated upwards within 50 s (trajectories shown in Fig. 6A, example in Video S7, panel A). This is demonstrated by the high proportion of larvae (3 batches, total *n*=100) moving in an upwards direction (Fig. 6B) in the seconds after mixing, in addition to a positive vertical displacement of 0.5 mm s⁻¹ (Fig. 6C). This indicates that our *A. millepora* larvae were positively buoyant in seawater at this developmental stage of 5 dpf.

**Figure 6.**
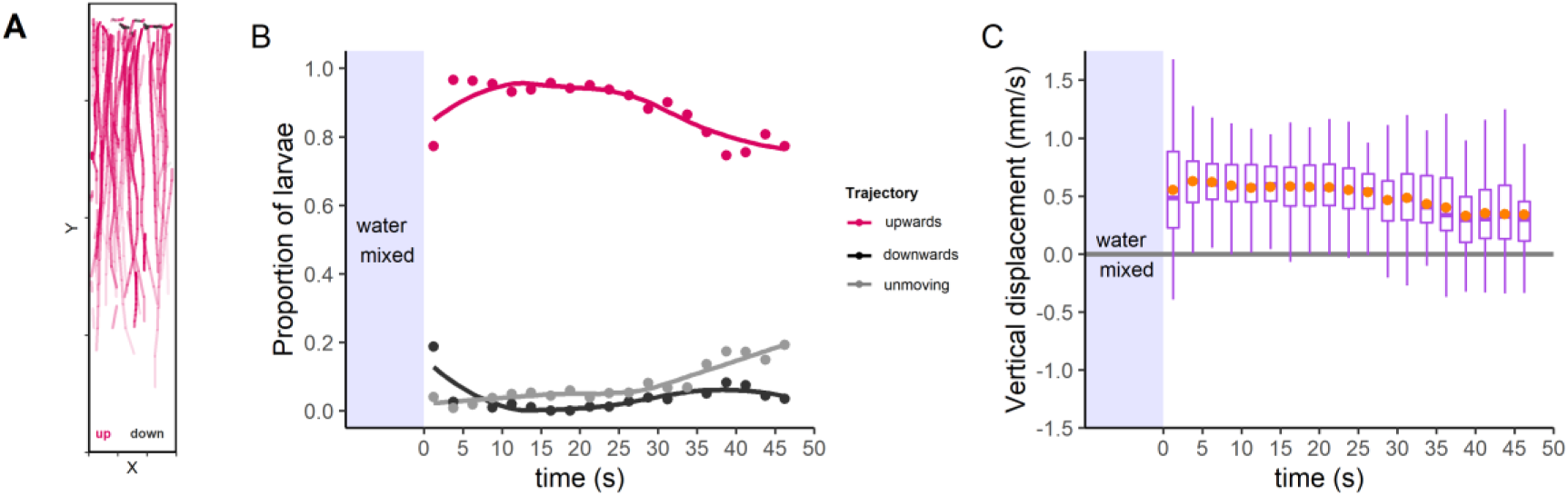
Vertical trajectories of larvae with swimming cilia removed. Data are averaged from three batches containing a total of n=96 tracked larvae. **A)** Example batch showing trajectories of larvae over time. Red lines show movement upwards in the cuvette whereas black shows downwards movement, and increasing time is shown by increasing weight (alpha value) of that line. **B)** Mean proportion of larvae moving upwards (red), downwards (black) or neither (grey) after mixing at each timepoint. Local regression curves fitted over points using LOESS-smoothing. **C)** Vertical displacement shown as boxplots with mean values overlaid (orange points). Boxes represent interquartile range with median centre line; whiskers extend to the most extreme values within 1.5 × the interquartile range. Positive values indicate net upward movement.

To test the hypothesis that the ciliary arrest and body rounding response to darkness results in a similar upwards movement of the larvae, we subjected them to 90 s on/off periods of bright white light and tracked their vertical position in the same cuvette. Larval swimming speed showed similar trends to our horizontal cuvette assay, where speed significantly reduced during the dark periods (Wilcoxon rank sum test, *W*=52652241, p<0.001) (Supplementary Table S4, Fig. S5).

In the first light-on period, the proportion of larvae swimming up and down were relatively similar as they explored the cuvette (Fig. 7) and this period was therefore excluded from statistical testing. However, when the light stimulus was turned off, over half of the larvae showed net upward movement in darkness. In the following light phase, the proportion of larvae moving in a downward direction became dominant (>0.5), before a return to upward movement in the final dark phase. It should be noted that the mean distances travelled were small and vertical displacement shows only a weak, but significant trend of upward movement in the dark, and downwards movement in the second light phase (Wilcoxon rank sum test, *W*=87668661, p<0.001). Batch 1 was an exception to this trend and showed no difference between light and dark phase vertical displacement (See supplementary Table S5 and Figs S6 & S7 for full results). Larvae closest to the bottom surface of the cuvette tended to exhibit a probing behaviour on the bottom in the light-on phase (Video S7, panel B). Then in the dark period, larvae in the bottom region became relatively still and floated very gradually upwards (Video S7, panel C). Larvae further up in the water column tended not to respond to the light dimming as often.

**Figure 7.**
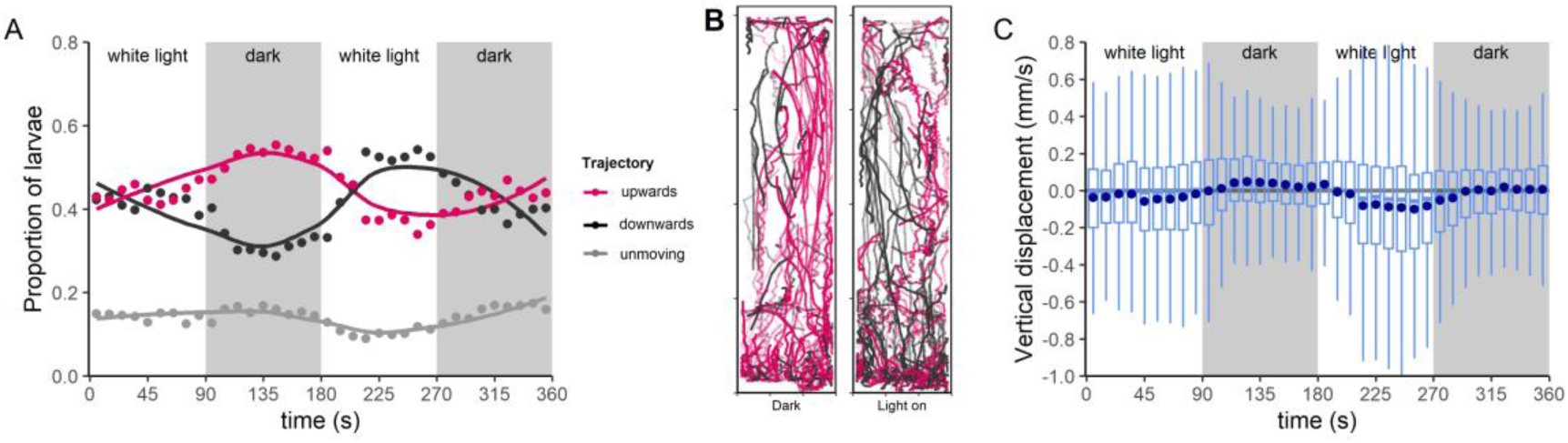
Vertical trajectories of larvae in the cuvette’s water column during on/off periods of bright white light and darkness. Data are averaged from six batches containing a total of n=361 tracked larvae. Grey shaded areas represent the two periods of darkness in time **A)** Mean proportion of larvae moving upwards (red), downwards (black) or neither (grey) at each timepoint. Local regression curves fitted over points using LOESS-smoothing. **B)** Example batch showing trajectories of larvae over time in the first dark period (91-180 s) and following light period (181-270 s). Red lines show movement upwards in the cuvette whereas black shows downwards movement, and increasing time is shown by increasing weight (alpha value) of that line. **C)** Vertical displacement as boxplots, with mean values overlaid (dark blue points). Boxes represent interquartile range with median centre line; whiskers extend to the most extreme values within 1.5 × the interquartile range. Positive values indicate net upward movement, whereas negative values indicate downward net movement.

Figure 7B shows an example batch (#2) with a proportionally greater number of smooth upward trajectories (red) indicating passive movement upwards in the dark, compared with the following light phase. Downward movement is dominant in the light phase, shown by the increased amount of black helical tracks, concentrated at the bottom of the cuvette. Note, the large amount of variation in general swimming behaviour, likely due to a portion of larvae who did not respond to the light dimming, retaining relatively fast movement and adding considerable noise to the data. The light-off phases were characterised by very slow and likely passive movement, with smaller distances travelled. The vertical displacement (Fig. 7C) does not reach the large positive values of deciliated larvae (Fig. 6C, see Video S7) and there is not a net migration to the top of the cuvette in the dark periods. This suggests that the cilia are perhaps important for maintaining position in the water column during periods of dark-induced contraction.

## Discussion

We demonstrate here that coral larvae of *A. millepora* exhibit a behavioural response to sudden light dimming by contracting the body and becoming rounded in shape (Video S2). They also reduce swimming speed, which is, at least partially caused by an arrest of the swimming cilia. We found no evidence of phototaxis, but larvae may be capable of directional swimming up and down in the vertical plane, using gravitaxis (Arai et al., 1993; Takeda-Sakazume et al., 2022). We interpret this light-off response as an important mechanism for maintaining position within suitable light envionments, allowing the larvae to further explore local habitats for settlement, or avoid entering unsuitable areas.

Our results for swimming speed reductions and lack of phototaxis behaviour were consistent with that for another acroporid coral larva, *A. tenuis* (Sakai et al., 2020). The maximum speed recorded from a single larva in the first bright light phase was 2.0 mm s⁻¹, suggesting that this species can swim at comparable speeds to *A. tenuis.* The mean free-swimming speed (0.41 ±0.43 mm s⁻¹) under bright white light was lower than that reported for *A. tenuis* (1.96 ±0.8 mm s⁻¹ (Sakai et al., 2020)), but this is mostly likely due to larger body size of *A. millepora* larvae, in addition to our use of a relatively smaller arena. Larvae made more frequent contact with its edges, unable to reach the speeds that they would otherwise achieve in larger water volumes. Tracking 3D movement of animals in a 2D plane also leads to extra underestimation of speed. However, despite the limitations of a confined laboratory experiment, the reduction in movement following light dimming was robust and reproducible across batches.

The muscular and ciliary light-off responses are robust, however not always consistent. Some individuals failed to respond during particular trials or did not respond at all. Internal state may contribute to this variability, as larvae were handled prior to experiments or tethered directly using suction micropipettes. Such handling may occasionally elicit stress or escape responses that override light-dependent modulation of swimming. In addition, the experimental arenas contained relatively high larval densities compared to natural reef environments, increasing the likelihood of collisions.

Following light dimming, a delay typically preceded both muscular contraction and ciliary arrest. This delay suggests that larvae may integrate light intensity over several seconds rather than responding instantaneously. Such temporal integration could allow larvae to filter out rapid, ecologically irrelevant fluctuations in light intensity, such as caustic flicker generated by surface waves (McFarland and Loew, 1983) or transient shadows cast by moving organisms. Short-lived reductions in light intensity, such as those encountered when swimming beneath small structures in the complex reef topography, may therefore not elicit a full locomotor response.

In *A. millepora* larvae, ciliary propulsion and muscular contraction appear to be two distinct motor systems that are engaged together during the light-off response. Ectodermal cilia generate metachronal waves that propel and rotate the larva through the water (Poon et al., 2023). In other ciliated larvae, such as *Platynereis*, ciliary arrest is coordinated by dedicated ciliomotor circuits (Marinković et al., 2020; Verasztó et al., 2017). Whether the diffuse nerve net of cnidarians contains comparable neural organisation remains unknown. Ciliary beating and particle image velocimetry measurements revealed that ciliary closures do not occur simultaneously across the body surface and there is spatial heterogeneity and temporal variability of ciliary arrest. This further supports a model in which the light-off response is mediated by slow, distributed signalling rather than fast, synchronised motor control. The relatively slow propagation rate of ciliary arrest and its spatial patchiness are consistent with a slow-acting signalling mechanism, potentially mediated by neuropeptides rather than fast synaptic transmission (Jékely and Yuste, 2024; Marinković et al., 2020). Notably, gentle persistent beating of cilia at the aboral pole, potentially associated with cilia of the apical organ, may contribute to the slow axial rotation observed in freely swimming larvae during dark periods, and suggests that their activity is controlled separately. This is a potential avenue for future research.

Muscular contraction similarly progressed gradually rather than as a synchronous whole-body event. We did not observe rapid, coordinated shortening across the oral–aboral axis. Instead, contraction appeared to occur in discrete events across different muscle groups (Video S6). This temporal pattern further supports the involvement of diffusible neuromodulatory signals rather than fast motor neuron activation (Jékely and Yuste, 2024). Muscular control of locomotion is thought to have evolved later than ciliary swimming in marine microswimmers (Jékely, 2011), raising the possibility that these two systems are only loosely coupled in coral larvae.

Opsin expression patterns indicate that photoreceptors in coral and other anthozoan planulae are distributed broadly across the ectoderm and endoderm rather than clustered into eye-like organs (Mason et al., 2012; McCulloch et al., 2023). Consequently, coral larvae lack the anatomical basis for spatial vision or directional light sensing (Brodrick and Jékely, 2023). Thus, larvae are unlikely to detect the direction of incoming light and instead respond to changes in overall irradiance. Simply put, the coral larva is unable to “see” where it is going when exploring the reef and cannot predict the light environment they are swimming toward.

Coral reefs are highly structured environments composed of complex three-dimensional structures formed by living and dead coral colonies (Guendulain-García et al., 2023).This architecture generates strong spatial heterogeneity in light intensity, with bright exposed surfaces interspersed with shaded crevices and overhangs. Light availability is a key factor underlying depth-related zonation of coral communities (Lesser et al., 2009; Roberts et al., 2019; Tamir et al., 2019), mediated by both larval settlement behaviour (Foster and Gilmour, 2016; Gleason et al., 2006; Mason et al., 2011; Mundy and Babcock, 1998; Ricardo et al., 2021; Strader et al., 2015) and post-settlement growth and survival (Hennige et al., 2010; Tamir et al., 2019; Vermeij and Bak, 2002). To locate an optimal settlement site in this habitat, larvae must use a variety of sensory cues, including light availability. Too shallow, and the colony would be exposed to harmful ultraviolet radiation and wave action (Lesser, 1997), and too deep or shaded would limit zooxanthellate symbiont recruitment and photosynthetic output. However, short-timescale swimming responses to light should not be conflated with long-term settlement preferences, which integrate multiple additional cues including substrate properties, biofilms, chemical signals, and hydrodynamic conditions.

Two possible interpretations of the light-off response are illustrated schematically in Fig. 8. One is that it limits the ingress of coral larvae into persistently dark microhabitats such as shaded crevices and overhangs (Fig 8,A). Sudden reductions in irradiance, such as those encountered when entering a shaded region of the reef, induce temporary slowing, body rounding, and intermittent axial rotation. Based on an estimated swimming speed of ∼1 mm s⁻¹ and a response delay of ∼15 s, a larva would travel only ∼15 mm into a shaded region before substantially reducing its movement, thereby limiting deeper penetration into darker spaces. The rounded body shape and gentle rotational behaviour may facilitate changes in swimming orientation, albeit in a non-directional manner, consistent with the absence of spatial light sensing in coral larvae (Fig. 8A). Random reorientation strategies are widely used by motile microorganisms, including bacteria and protists performing random-walk chemotaxis (Wan and Jékely, 2021), and a comparable mechanism in coral larvae could allow coarse regulation of position with respect to local light conditions. During prolonged periods of darkness, however, *A. millepora* larvae eventually elongate and resume exploratory swimming, as also reported for *A. tenuis* (Sakai et al., 2020). Therefore, it may serve to prevent irreversible confinement within shaded regions and allow the eyeless coral larva to navigate back towards more optimal light environments.

**Figure 8.**
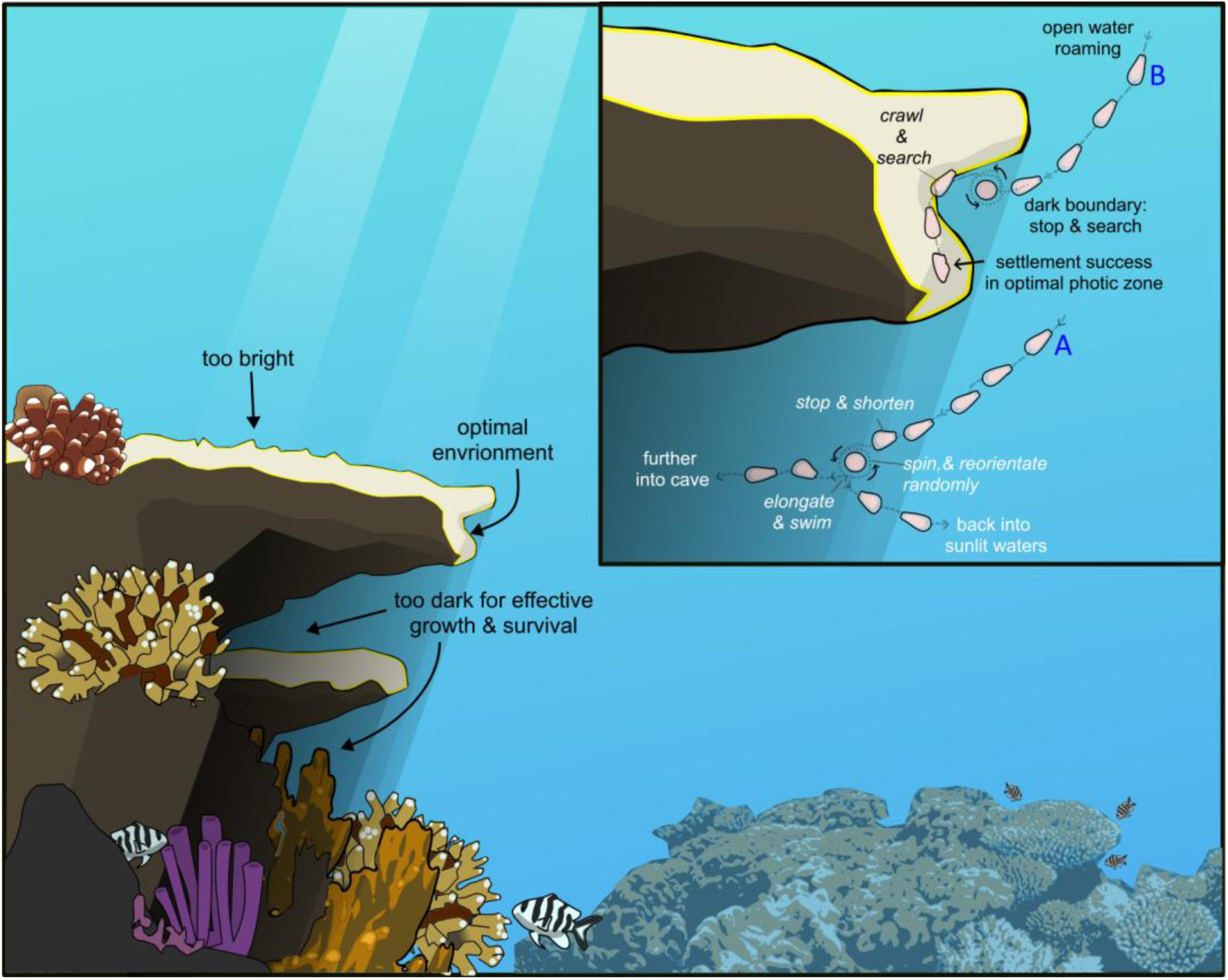
Conceptual schematic illustrating two non-exclusive possible interpretations of the larval light-off response in a coral reef habitat. Complex reef structure creates sharp light gradients between illuminated open water, partially shaded surfaces, and persistently dark microhabitats. **Insert: A-** The light-off response may limit further ingress into persistently dark regions by inducing temporary slowing, rounding, and random reorientation after sudden light dimming. **B-** The response may increase local sampling at light–dark boundaries by triggering a switch from rapid swimming to a localised exploratory behavioural state, consistent with an area-restricted search strategy.

An alternative but not mutually exclusive interpretation is that the light-off response modulates larval sampling behaviour at boundaries between illuminated and shaded regions of the reef (Fig. 8-B). During exploratory swimming in open, well-lit water, larvae swim actively and cover relatively large distances. A sudden reduction in irradiance may therefore act as a contextual cue indicating proximity to reef structure, triggering a rapid transition from active exploration to a localised behavioural state characterised by slowing, rounding, and intermittent rotation. This behavioural switch is consistent with the concept of “area-restricted search”, in which organisms adaptively transition between extensive and intensive movement modes in response to ecologically relevant cues, without requiring spatial vision or explicit target gradients (Dorfman et al., 2022). In the vertical cuvette experiments (Video S7), larvae that remained close to the bottom surface (where probing behaviour was frequently observed) responded most consistently to light dimming by reducing swimming and floating passively. In contrast, larvae higher in the water column often continued active swimming during dark periods. This context-dependence suggests that the light-off response may be activated by proximity to surfaces, consistent with active habitat exploration rather than pelagic dispersal. By reducing forward movement at light–dark boundaries, larvae may retain position near the interface between open water and shaded reef surfaces, rather than passing by quickly. These partially illuminated surfaces may meet the requirements for their photic zone preference, and such behaviour could further increase sampling time of local sensory cues (e.g. chemical signals for algae and biofilms) at microhabitats that serve as potential settlement sites. Importantly, this interpretation does not require larvae to prefer or move toward darker regions, but instead suggests that light dimming serves as a trigger for behavioural switching when larvae encounter the structural complexity of the reef. Therefore, we propose that the light-off response functions as a mechanism for detecting and responding to light-dark boundaries, rather than as an indicator of preference for either light or darkness.

Taken together, these hypotheses suggest that the light-off response may influence both large-scale positioning within the reef light-landscape and fine-scale interactions with potential settlement sites. To our knowledge, this is the first report of a body-rounding response to light dimming in coral larvae. Our study raises several avenues for future investigation, including how larvae respond to gradual rather than abrupt changes in irradiance along depth-light gradients, and how neural and biomechanical coupling between ciliary and muscular systems is achieved.

## Acknowledgements

The research presented in this manuscript was conducted at the Living Systems Institute, University of Exeter. It was funded by the Human Frontiers Science Program (HFSP) grant No. RGP0033/2020. This project has also received funding from the European Research Council (ERC) under the European Union’s Horizon 2020 research and innovation programme (Grant Agreement No. 101020792 to GJ and No. 853560 to KYW).

We would like to thank the staff of the Horniman Museum and Gardens aquarium facility and the Coral Spawning Lab for providing us with live coral larvae. Thank you also to Prof. Nick Roberts from the Ecology of Vision Lab, Bristol for lending us a spectrometer.

## Supplementary material

**Figure S1.**
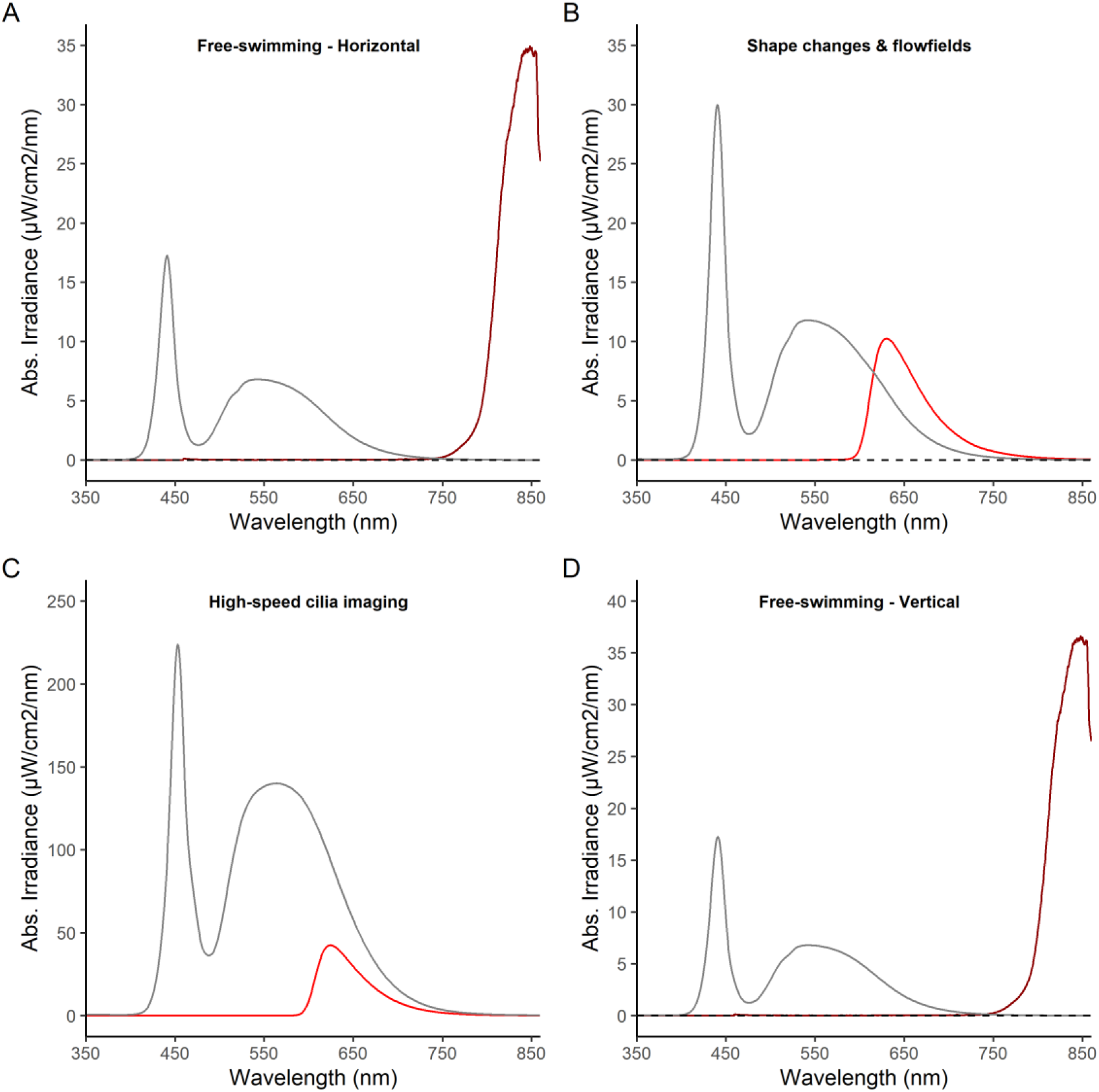
Absolute irradiance (µW/cm2/nm) light spectra measured from experimental apparatus. **A)** Free-swimming experiments using the shallow cuvette to assess horizontal movement. Bright white LED stimulus when on (grey solid line) and off (black dashed line). Solid dark-red line shows the near-infrared ring light, used constantly for camera illumination. **B)** Tethered larvae experiments to assess larval shape changes and flow fields. Bright white LED stimulus when on (grey solid line) and off (black dashed line). Solid red line shows the dim red-filtered illumination used for imaging. **C)** High-speed imaging of tethered larvae to assess ciliary beating. Bright white light stimulus when on (grey line) and when passed through a red filter (red line). **D)** Free-swimming experiments using the tall cuvette to assess vertical movement. Bright white LED stimulus when on (grey solid line) and off (black dashed line). Solid dark-red line shows the near-infrared ring light, used constantly for camera illumination. All spectra measurements were made from the position of the experimental cuvette where the larvae were held.

**Table S1.** Wilcoxon rank sum statistical tests (W) comparing swimming speed (mm s⁻¹) in the second half of dark and light time periods for the horizontal free-swimming experiments. Columns 2 and 3 are median swimming speed values for each light condition. The bottom row shows the test result when data from all batches were pooled together.

| batch | dark_med | Light-on_med | W-value | p-value | signif. |
| --- | --- | --- | --- | --- | --- |
| 1 | 0.050 | 0.340 | 3062 | $6.80 \text{ e-}15 < \text{s}^{-1}\text{up} >$ | *** |
| 2 | 0.039 | 0.241 | 4141 | $8.75 \text{ e-}27$ | *** |
| 3 | 0.039 | 0.372 | 2916 | $1.55 \text{ e-}18$ | *** |
| 4 | 0.044 | 0.151 | 4618 | $1.02 \text{ e-}11$ | *** |
| 5 | 0.083 | 0.212 | 6730 | $8.89 \text{ e-}08$ | *** |
| 6 | 0.123 | 0.241 | 8905 | 0.0218 | * |
| **pooled** | **0.058** | **0.235** | **183558** | **2.91 e-63** | **\*\*\*** |

**Figure S2.**
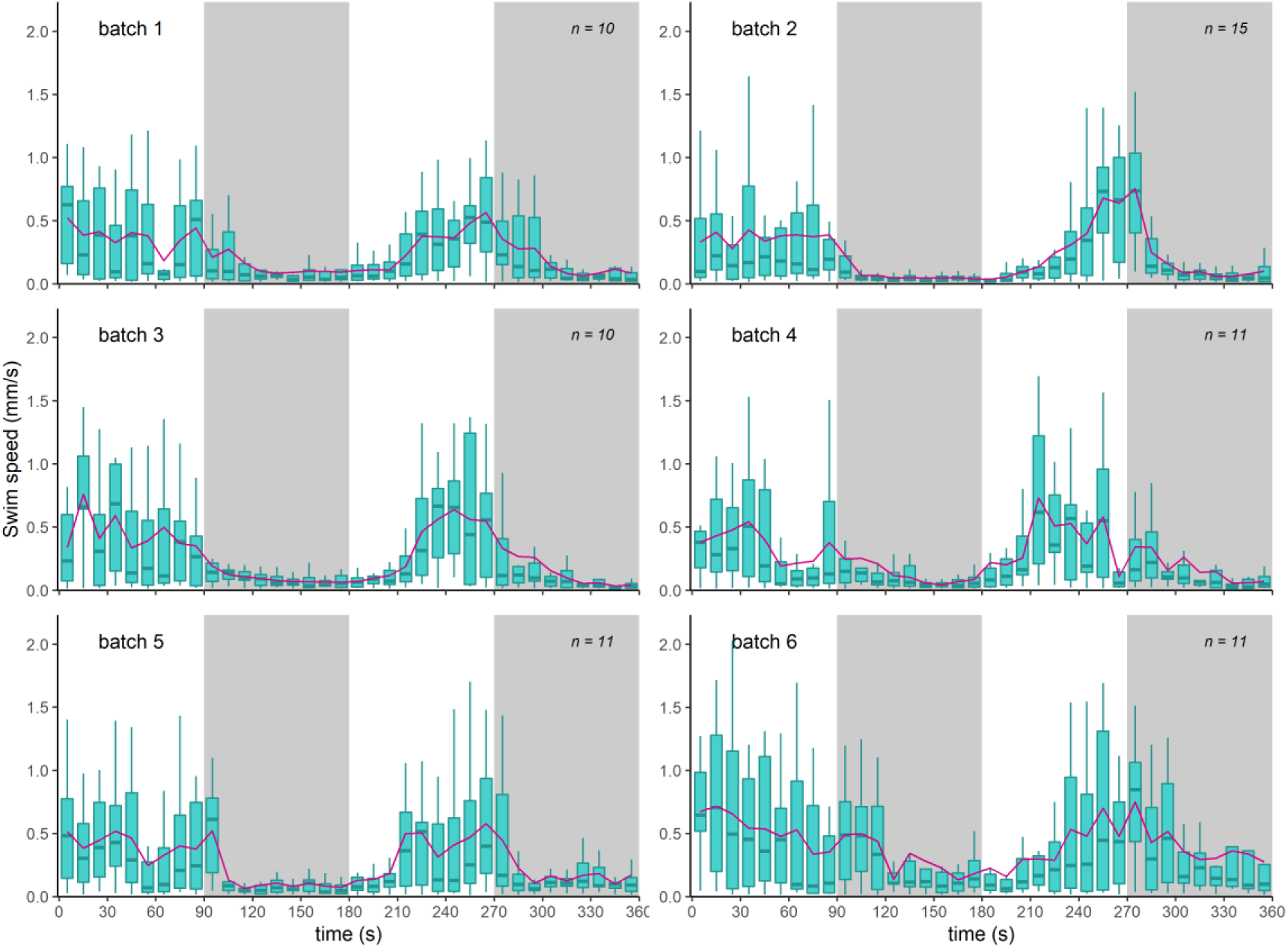
Swimming speed data (mm s⁻¹) for six batches of larvae in the horizontal free-swimming cuvette experiment. Data are represented as boxplots of the interquartile range with median line and whiskers extending to the most extreme values within 1.5× the interquartile range. Purple line represents mean swimming speed. White areas represent time periods with the light stimulus being on, whereas shaded boxes show the dark time periods.

**Table S2.** Wilcoxon rank sum statistical tests (W) comparing horizontal displacement (mm s⁻¹) in the second half of dark and light time periods for the horizontal free-swimming experiments. Positive median values in columns 2 and 3 represent movement toward the light source on one side and negative values represent movement away from it. The bottom row shows the test result when data from all batches were pooled together.

| batch | dark_med | light-on_med | W-value | p-value | signif. |
| --- | --- | --- | --- | --- | --- |
| 1 | 0.019 | -0.004 | 6878 | 0.4424 | - |
| 2 | -0.064 | 0.140 | 10995 | 0.0081 | ** |
| 3 | -0.016 | 0.008 | 8234 | 0.7845 | - |
| 4 | -0.004 | -0.005 | 8397 | 0.4157 | - |
| 5 | 0.015 | -0.038 | 11484 | 0.2054 | - |
| 6 | -0.044 | -0.066 | 10555 | 0.9939 | - |
| **pooled** | **-0.023** | **-0.004** | **338476** | **0.2727** | **-** |

**Figure S3.**
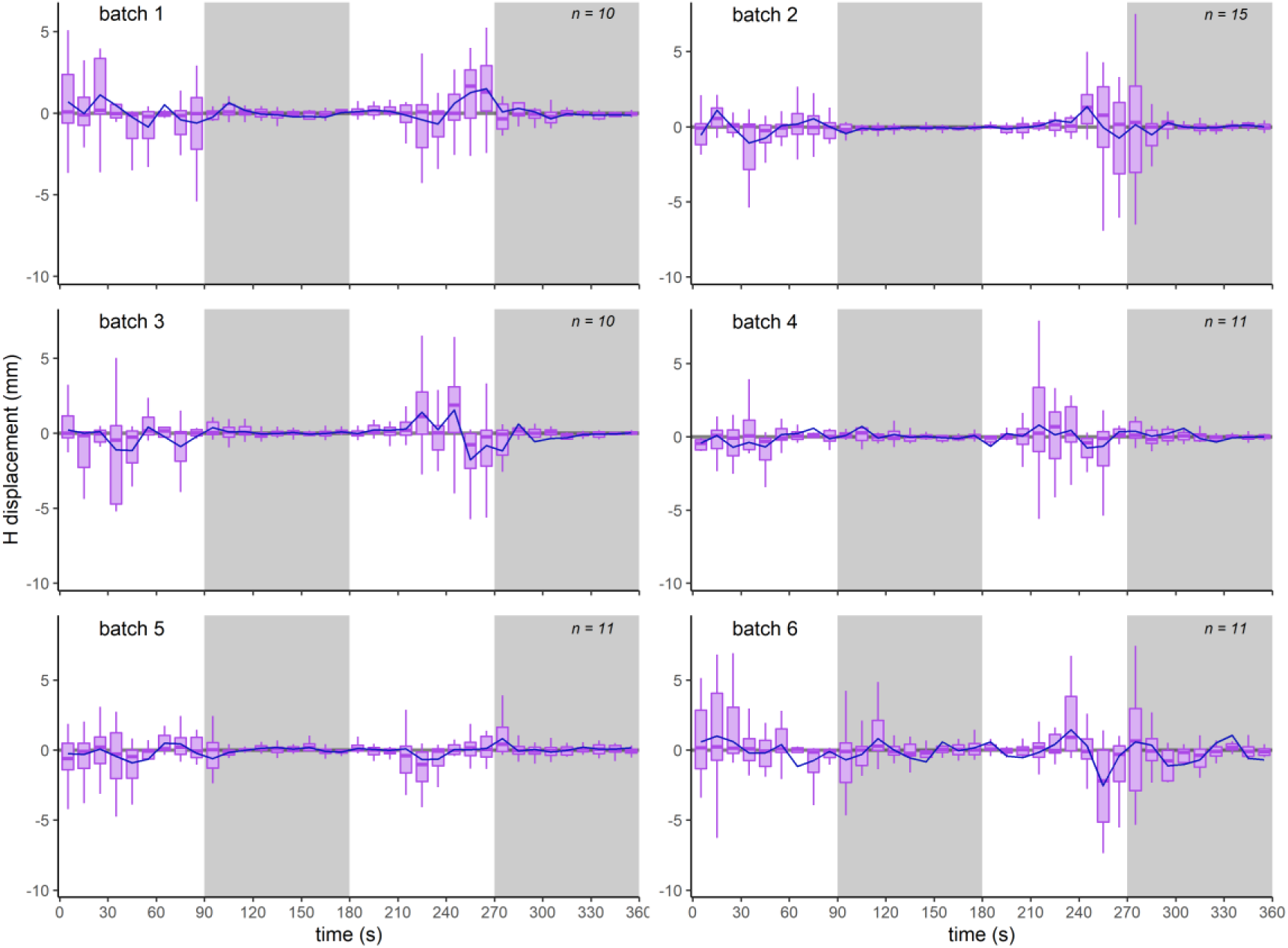
Horizontal displacement data (mm s⁻¹) for six batches of larvae in the horizontal free-swimming cuvette experiment. Data are represented as boxplots of the interquartile range with median line and whiskers extending to the most extreme values within 1.5× the interquartile range. Dark blue line represents mean swimming speed. White areas represent time periods with the light stimulus being on, whereas shaded boxes show the dark time periods.

**Figure S4.**
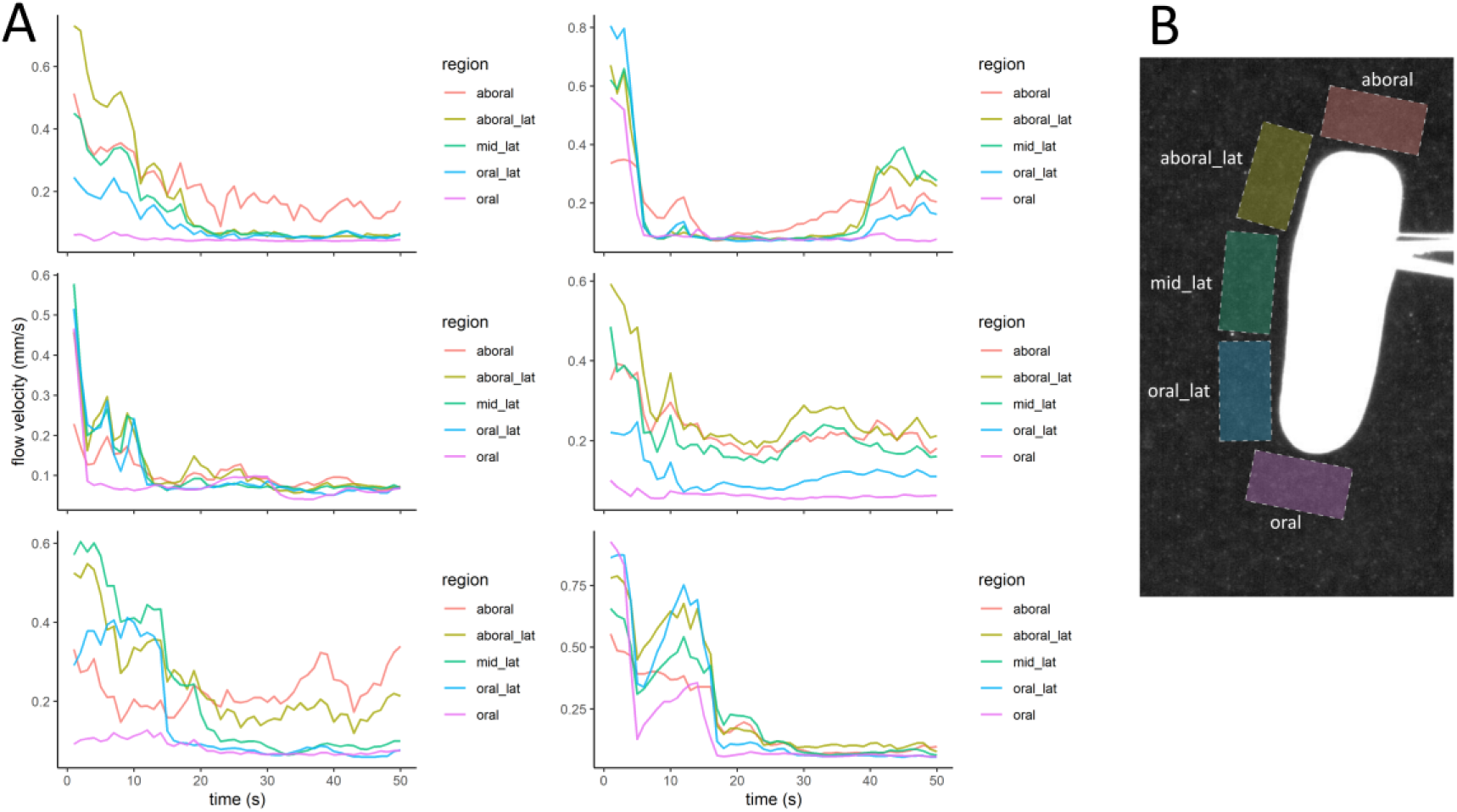
Spatially resolved flow velocities measured by particle image velocimetry (PIV) around tethered larvae. **(A)** Flow velocity (mm s⁻¹) within five regions spanning the aboral to oral pole is shown for each larva (one plot per individual). Colours indicate the sampled regions. Time 0 corresponds to the moment when LED illumination was switched off, marking the start of the “dark” period. The first 50 s of this 90 s dark interval are shown to illustrate the light-off response. **(B)** Representative example showing the five spatial regions from which flow velocity was quantified using PIV.

**Table S3.** Wilcoxon rank sum statistical tests (W) comparing the aspect ratio (length/width) and 2D areas of larval profiles in the second (45 s) half of dark and light time periods. Median values are shown for the light periods for both 2021 and 2022 batches of larvae.

| year | dark_med | light_on_med | W-value | p-value | signif. |
| --- | --- | --- | --- | --- | --- |
| **Aspect ratio** |  |  |  |  |  |
| 2021 | 1.35 | 1.51 | 1058.5 | 0.0014 | ** |
| 2022 | 1.22 | 2.15 | 551 | 8.50 e-11 | *** |
| **2D area** |  |  |  |  |  |
| 2021 | 234 | 256 | 1262 | 0.0425 | * |
| 2022 | 215 | 243 | 716 | 1.89 e-08 | *** |

**Figure S5.**
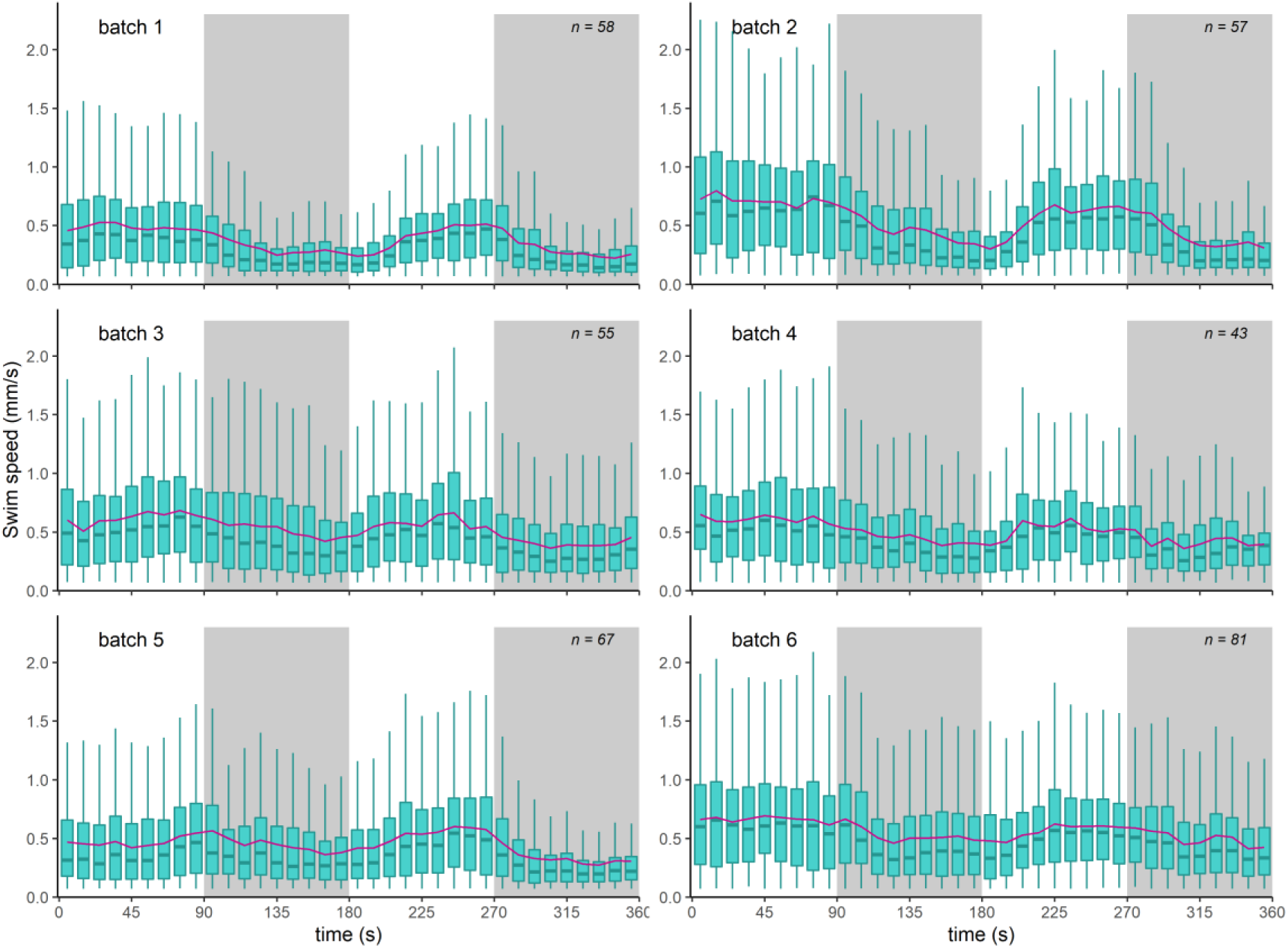
Swimming speed data (mm s⁻¹) for six batches of larvae in the vertical free-swimming cuvette experiment. Data are represented as boxplots of the interquartile range with median line and whiskers extending to the most extreme values within 1.5× the interquartile range. Purple line represents mean swimming speed. White areas represent time periods with the light stimulus being on, whereas shaded boxes show the dark time periods.

**Table S4.** Wilcoxon rank sum statistical tests (W) comparing swimming speed (mm s⁻¹) in the middle 30 sec of both dark and the second light periods for the vertical cuvette free-swimming experiments. Columns 2 and 3 are median swimming speed values for each light condition. The bottom row shows the test result when data from all batches were pooled together.

| batch | dark_med | light_on_med | W-value | p-value | signif. |
| --- | --- | --- | --- | --- | --- |
| 1 | 0.175 | 0.375 | 1396172 | 2.59 e-119 | *** |
| 2 | 0.252 | 0.535 | 982012 | 9.15 e-71 | *** |
| 3 | 0.313 | 0.531 | 1527658 | 1.43 e-36 | *** |
| 4 | 0.325 | 0.537 | 754550 | 2.89 e-31 | *** |
| 5 | 0.248 | 0.442 | 1753602 | 2.68 e-46 | *** |
| 6 | 0.355 | 0.537 | 2545960 | 2.67 e-38 | *** |
| **pooled** | **0.266** | **0.482** | **52652240.5** | **1.47 e-296** | **pooled** |

**Figure S6.**
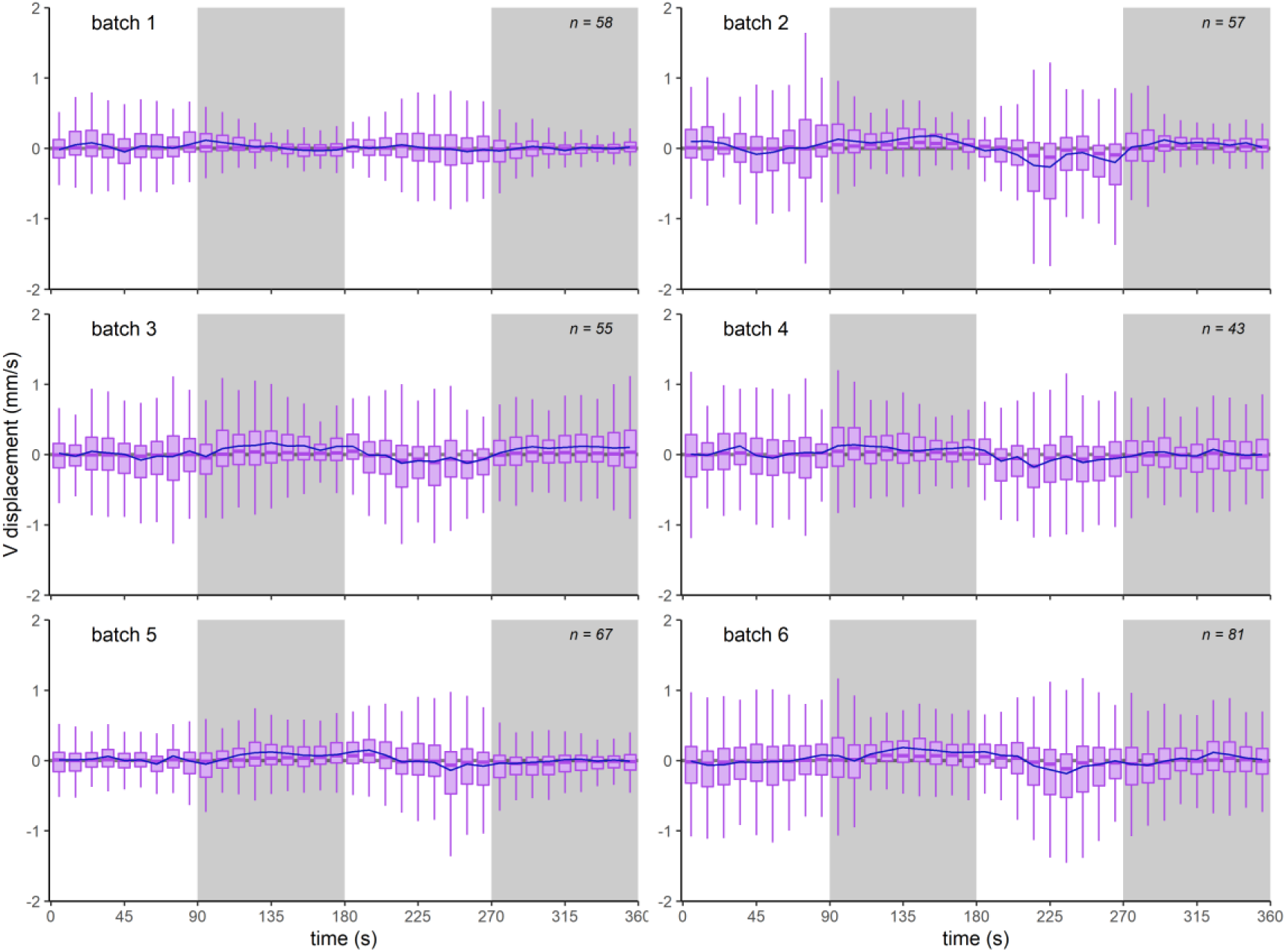
Vertical displacement data (mm s⁻¹) for six batches of larvae in the vertical free-swimming cuvette experiment. Data are represented as boxplots of the interquartile range with median line and whiskers extending to the most extreme values within 1.5× the interquartile range. Dark blue line represents mean swimming speed. White areas represent time periods with the light stimulus being on, whereas shaded boxes show the dark time periods.

**Table S5.** Wilcoxon rank sum statistical tests (W) comparing vertical displacement (mm s⁻¹) in middle 30 sec of both dark and the second light periods for the vertical cuvette free-swimming experiments. Positive median values in columns 2 and 3 represent movement toward the light source on one side and negative values represent movement away from it. The bottom row shows the test result when data from all batches were pooled together.

| batch | dark_med | light_on_med | W-value | p-value | signif. |
| --- | --- | --- | --- | --- | --- |
| 1 | 0.011 | 0.011 | 2325504 | 0.170 | - |
| 2 | 0.060 | -0.091 | 2118636 | 1.48 e-76 | *** |
| 3 | 0.027 | -0.085 | 2563379 | 1.91 e-51 | *** |
| 4 | 0.019 | -0.067 | 1221317 | 9.83 e-20 | *** |
| 5 | 0.007 | -0.010 | 2524931 | 6.66 e-05 | *** |
| 6 | 0.042 | -0.063 | 4121012 | 5.28 e-62 | *** |
| **pooled** | **0.025** | **-0.040** | **87668662** | **3.29 e-151** | **\*\*\*** |

**Figure S7.**
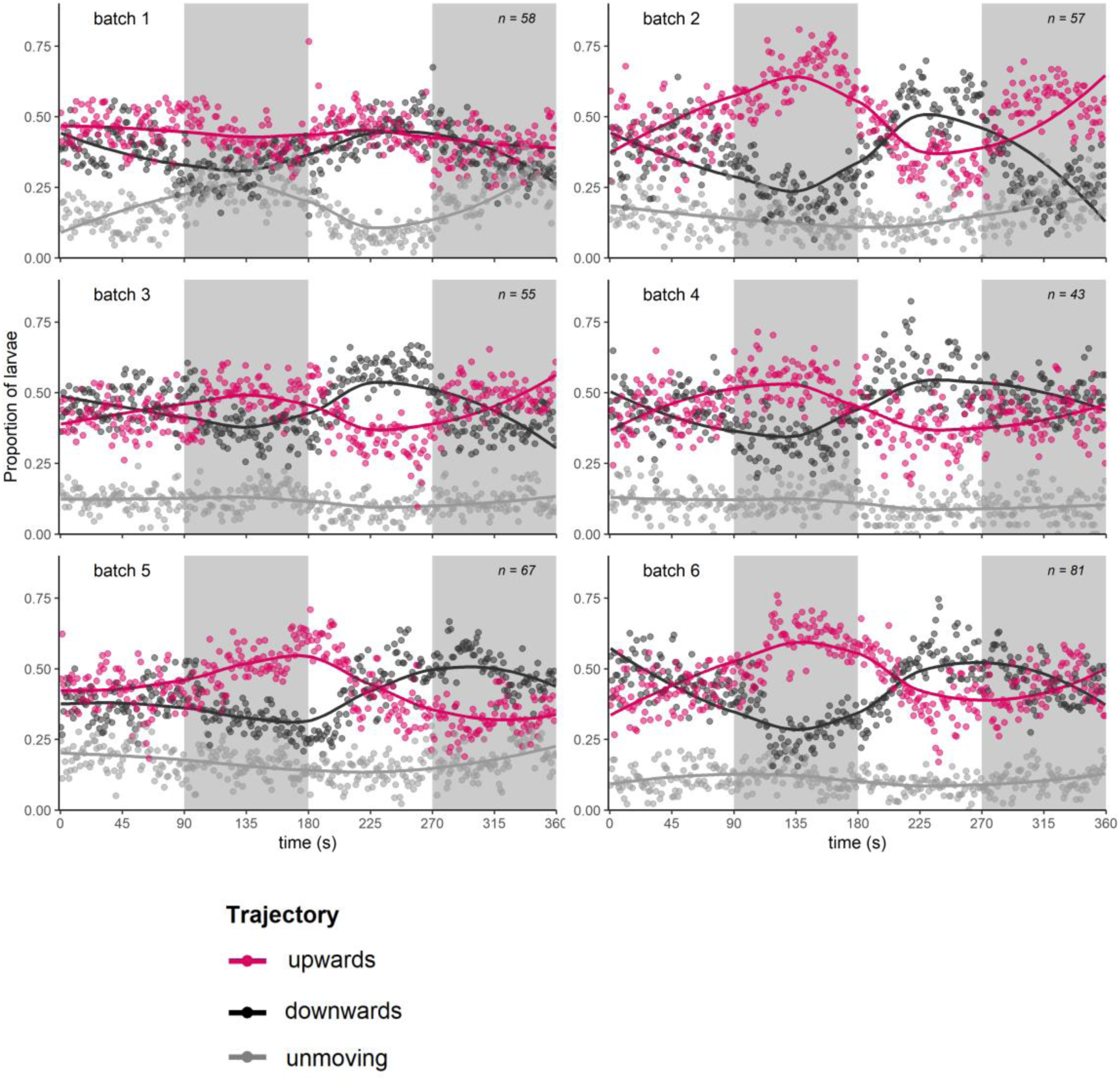
Swimming trajectory data for six batches of larvae in the vertical free-swimming cuvette experiment. Data are represented as the proportion of total larvae at each time point that travelled upwards (red), downwards (black) or neither (grey) at each timepoint. Local regression curves fitted over points using LOESS-smoothing.. White areas represent time periods with the light stimulus being on, whereas shaded boxes show the dark time periods.

## Video legends

Video data supporting this manuscript are available online at: (Brodrick et al., 2025).

**Video S1. Horizontal tracking of coral larva trajectories in shallow cuvette.** The video is split into four panels to show each of the 90 s light-on and dark phases. Playback is sped up by 10x. Tracks are coloured by larva ID and persist for each light phase showing the total trajectory. Light-on periods are characterized by faster swimming, covering greater distances in the cuvette. During dark periods, the larvae often tended to spin on the spot.

**Video S2. Tethered larva performing its light-off response to sudden intensity dimming.** The larvais held in seawater containing nanoparticle beads to allow measurement of flow speeds via PIV. The first 5 s of the video show the larva elongated with beating ectodermal cilia under bright white light. At 5 s, the bright light is dimmed leaving the larva in dim red illumination only, which prompts the response of contracting the body into a round ball, and later, all flow stops around the larva as a result of ciliary arrest. Playback is in real time.

**Video S3. Particle image velocimetry (PIV) analysis.** Example of a tethered larva (shaded black) with flow velocity in the surrounding seawater represented by arrows and colour (scale on left). The white light stimulus is on when the video starts but is dimmed 5 s later. All flow stops in the water surrounding the larva around 15 s after light dimming, suggesting ciliary arrest.

**Video S4. Ectodermal ciliary arrest of larva #1 in response to white light dimming.** The video begins with white light (WL) on, then at 33.4 s, the white light is dimmed, and imaging continues using dim red illumination only (RL). An arrest of beating can be seen among the cilia in focus 4 s later (at 38 s). Video playback is in real time, but it has been compressed (from 500 fps) to 25 fps.

**Video S5. Ectodermal ciliary arrest of larva #2 in response to white light dimming.** The video begins with white light (WL) on, then at 33 s, the white light is dimmed, and imaging continues using dim red illumination only (RL). An arrest of beating can be seen spreading down through the cilia in focus, starting at 77 s (43 s after the light dimming). Video playback is in real time, but it has been compressed (from 500 fps) to 25 fps.

**Video S6. Coral larvae contract the body in response to light dimming.** Two examples of coral larvae are shown from cohorts from consecutive spawning years. The larvae are tethered on a suction micropipette as the light stimulus is turned on and off, twice. Video playback is 10x. Note the muscle contraction events in dark periods, gradually rounding the body shape. An elongated shape is resumed in bright light periods via muscle relaxation.

**Video S7. Vertical tracking of larval trajectories in tall cuvette.** Fading trajectory tails span 30 frames and are coloured according to mean track speed (blue represents slow travel, while red is fast). Video playback is sped up by 10x. **A)** A batch of deciliated and immobile larvae float upwards over 50 s after mixing to disperse them in the column. **B)** Larvae swimming actively with white light on. Magenta box in the second repeat of the video highlights the larvae probing the base of the cuvette. **C)** Same batch of larvae as in B, in darkness after the sudden dimming of white light. Yellow box in second video repeat highlights that most larvae near the bottom surface showed alight-off response and floated very slightly upward. Whereas larvae higher inthe column tended to continue active swimming as before.

